# Discovery of fungal-specific targets and inhibitors using chemical phenotyping of pathogenic spore germination

**DOI:** 10.1101/2021.02.06.430071

**Authors:** Sébastien C. Ortiz, Mingwei Huang, Christina M. Hull

## Abstract

There is a critical need for new antifungal drugs; however, the lack of available fungal-specific targets is a major hurdle in the development of antifungal therapeutics. Spore germination is a differentiation process absent in humans that could harbor uncharacterized fungal-specific targets. To capitalize on this possibility, we developed novel phenotypic assays to identify and characterize inhibitors of spore germination of the human fungal pathogen *Cryptococcus*. Using these assays, we carried out a high throughput screen of ~75,000 drug-like small molecules and identified and characterized 191 novel inhibitors of spore germination, many of which also inhibited yeast replication and demonstrated low cytotoxicity against mammalian cells. Using an automated, microscopy-based, quantitative germination assay (QGA), we discovered that germinating spore populations can exhibit unique phenotypes in response to chemical inhibitors. Through the characterization of these spore population dynamics in the presence of the newly identified inhibitors, we classified 6 distinct phenotypes based on differences in germination synchronicity, germination rates, and overall population behavior. Similar chemical phenotypes were induced by inhibitors that targeted the same cellular function or had shared substructures. Leveraging these features, we used QGAs to identify outliers among compounds that fell into similar structural groups and thus refined relevant structural moieties, facilitating target identification. This approach led to the identification of complex II of the electron transport chain as the putative target of a promising structural cluster of germination inhibitory compounds. These inhibitors showed high potency against *Cryptococcus* spore germination, while maintaining low cytotoxicity against mammalian cells, making them prime candidates for development into novel antifungal therapeutics.

## Introduction

Human fungal pathogens are an unmitigated problem causing ~1.5 million deaths a year worldwide (1). One of the biggest hurdles in the treatment of invasive fungal diseases is the lack of available therapeutics. There are three primary classes of antifungal drugs, all of which are suboptimal due to properties ranging from high toxicity to humans to rapid microbial resistance development (2–5). These classes target cell membranes or cell wall components, which have been the canonical targets for antifungal development (2,6). While these cellular structures provide fungal-specific targets, the deficiency of novel antifungal agents indicates that new fungal-specific targets need to be identified and exploited. However, due to the eukaryotic nature of fungi and resulting conservation of molecular moieties between humans and fungi, the identification of fungal-specific pathways has been difficult.

One proposed solution is to target the process of spore germination (7). Spores are dormant, stress-resistant cell types formed by many organisms to survive harsh environmental conditions and/or spread to new environments, and spores are infectious particles for most invasive human fungal pathogens (8,9). To cause disease, fungal spores must escape dormancy through the process of germination, a process that appears unlike any in humans, and grow vegetatively in the host. Due to its specialized nature, spore germination may involve fungal pathways distinct from those in humans. We hypothesized that the process of spore germination would harbor new fungal-specific targets, and compounds that inhibit germination would therefore be less toxic to mammalian cells. Thus, germination inhibitors would be prime candidates for development into antifungal drugs for the prevention and/or treatment of many invasive fungal diseases.

The development of new antifungal drugs has been slow relative to other antimicrobial agents such as those against bacteria and viruses, despite significant screening efforts (10). Traditionally, the primary method for identifying antifungal compounds was based on tracking changes in a biologically relevant readout such as fungal growth (Phenotypic Drug Discovery). While this approach yielded many antifungal compounds over the years, most were also toxic to mammalian cells. As molecular techniques advanced, researchers in many fields moved toward Targeted Drug Discovery, which relies on identification of inhibitors of a specific, known molecular target. This approach proved to be useful in some arenas; however, it largely failed for antifungal drug development presumably because of a lack of identified fungal-specific targets. As a result, no new classes of antifungal therapeutics have come to market in the last 20 years, and there is renewed interest in phenotypic drug discovery (2,11). However, because growth inhibition screening alone has been used exhaustively as a target phenotype in fungi, for the promise of phenotypic drug discovery to be fully realized, the field must improve upon traditional growth screening methods and/or couple them with novel assays (10).

To address this need, we used fundamental biological discoveries of pathogenic spore biology to drive the development of two new phenotypic assays that use spore germination as a readout. The first is a luciferase-based assay for high throughput screening of compounds to identify germination inhibitors, and the second is an automated, quantitative, microscopy-based germination assay for high-resolution evaluation of large populations of germinating spores. We developed these tools for use with the spores of the invasive human fungal pathogen *Cryptococcus*. This environmental budding yeast is the leading cause of fatal fungal disease worldwide, causing several hundred thousand deaths per year, particularly among people with compromised immune systems (5). The *Cryptococcus* system is known among human fungal pathogens to be well-developed with many molecular and genetic tools. In addition, *Cryptococcus* spores germinate synchronously under nutritionally favorable conditions and do so largely independent of spore density (12). These unique properties facilitated the development of the phenotypic germination assays that we used to identify and characterize 191 novel fungal germination inhibitors. We discovered that population level dynamics could be used to classify distinct chemical phenotypes, and we used those classifications to show that compounds with similar substructures demonstrated similar chemical phenotypes. This process led to the rapid identification of phenotypic outliers and facilitated target identification, resulting in the discovery of a novel set of fungal-specific electron transport chain inhibitors that are prime candidates for development into a new class of antifungal drugs for use in the prevention of fatal fungal diseases.

## Results

### Eukaryotic translation inhibitors prevent initiation of Cryptococcus spore germination

Prior studies of *Cryptococcus* spore germination showed that different conditions, mutants, and drugs alter the behavior of spore populations during germination (7,12). Based on these findings, we hypothesized that characterizing the behaviors of spore populations under different conditions would facilitate the identification of specific cellular processes required for spore germination. Because new protein synthesis is known to be required for successful germination in many fungi (13), we treated populations of spores under germinating conditions with the eukaryotic ribosome inhibitor cycloheximide, and evaluated their responses using our quantitative germination assay (QGA).

In this automated, microscopy-based assay, germination progression of spores is monitored as a function of changes in cell morphology over time (spores are small and oval; yeast are large and circular). Individual spores in a population (~1×10^4^ per sample) are measured to determine size (area) and shape (aspect ratio) from the onset of germination, and the data are collected for each cell over the time of germination (**Figure 1A**). Using the QGA, we determined that a concentration of 10 μM cycloheximide fully prevented spores from initiating any changes in size or shape, indicating full inhibition of germination and suggesting that new protein synthesis is required for *Cryptococcus* spores to germinate (**Figure 1B**).

**Figure 1.**
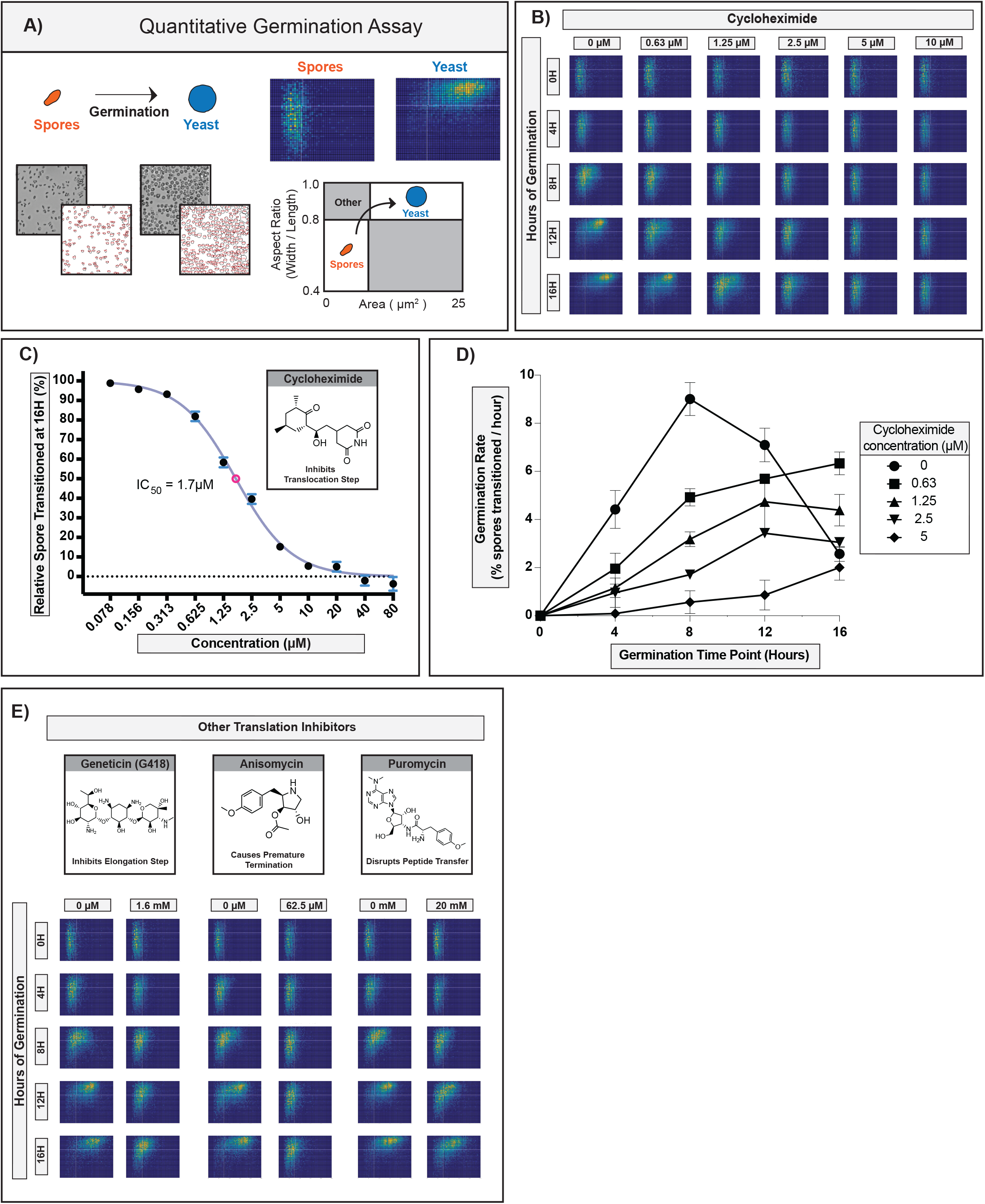
Evaluation of Eukaryotic Translation inhibitors in the QGA assay. A) Diagram of QGA assay, these data are represented in two-dimensional histograms that show the numbers of cells in each position as a function of pixel intensity. Spores (small, oval cells) populate the lower left quadrant, and yeast (large, circular cells) populate the upper right quadrant. B) Germination profiles of spores in the presence of 0, 0.63, 1.25, 2.5, 5, or 10 μM cyloheximide each plot representing ~6,000 spores. C) Dose response curve of spores transitioned at 16 hours relative to control in the presence of 0.078 to 80 μM cycloheximide (error bars represent standard deviation) D) Germination rates of spores at different time points in the presence of 0, 0.63, 1.25, 2.5, or 5 μM cyloheximide (error bars represent standard deviation). E) Germination profiles of translation inhibitors each plot representing ~6,000 spores.

As the concentration of cycloheximide decreased, the amount of germination increased, exhibiting concentration-dependent inhibition of germination with an IC_50_ of 1.7 μM (**Figure 1C**). In addition, we observed that even as the germination of the population of spores was slowing down due to inhibition, all of the spores responded in a similar manner and maintained their synchronous response (**Figure 1B**). While the overall rate of germination for the population changed in response to cycloheximide, other properties were unchanged (population synchronicity, pattern of morphological changes, integrity of individual spores), which facilitated the determination of specific rates of germination (i.e. transition out of the spore state) at each concentration tested (**Figure 1D**). We observed that as the concentration of cycloheximide increased, germination rates decreased, causing a “slow down” phenotype across the population.

From these data we concluded that new protein synthesis was likely required very early in the germination process. We further surmised that if the cycloheximide phenotype were specific to inhibition of protein translation (as opposed to off-target effects), other inhibitors of eukaryotic protein translation would produce the same phenotype. To test this hypothesis, we evaluated 3 structurally distinct inhibitors of eukaryotic protein translation (geneticin, anisomycin, and puromycin) and determined their effects in QGAs (**Figure 1E**). Although the inhibitors showed different potencies against germination (i.e. different concentrations were required to achieve similar effects), all three inhibitors caused a “slow down” phenotype that maintained population synchronicity, mimicking cycloheximide.

Together, these data show that for each inhibitor, the QGA was an effective method for determining a concentration-dependent phenotype, determining precise inhibitory concentrations, and quantitating changes in germination rates. Furthermore, the consistent phenotypes across translation inhibitors suggested that inhibitors targeting the same cellular function generate a similar phenotype in the QGA. These findings indicated that QGAs would be a powerful tool in the validation, prioritization, and characterization of diverse germination inhibitors with unknown targets.

### Combined HTS and QGA analysis identified 191 novel germination inhibitors

To identify potential inhibitors of spore germination, a NanoLuciferase (NL)-based high throughput screening assay was developed, and compounds from three libraries of structurally diverse, drug-like small molecules (LifeChem 1-3) were screened (**Figure S1)**. For the assay, a previously identified protein (CNK01510) was fused to the NanoLuciferase protein (Promega) and introduced into its endogenous locus in the *Cryptococcus* genome (14). Strains harboring the integrated NL protein fusion were crossed under sexual development conditions to produce spores. Spores from the NL strains yielded very little NL enzyme activity; however, yeast from those strains produced robust NL signal. Most importantly, the amount of NL signal correlated with germination state. As spores germinated into yeast, the levels of NL signal increased, resulting in a robust signal over baseline at the end of germination (~14-fold) (**Figure S1**). Inhibitors of germination (e.g. cycloheximide) caused low levels of NL signal, and solvent-only controls (e.g. DMSO) affected neither germination nor NL signal (**Figure S1**). Of the ~75,000 compounds screened, ~2100 compounds caused a ≥20% decrease in NL signal relative to the solvent-only control and were rescreened in duplicate. Compounds that showed ≥50% inhibition of the NL control signal were selected, resulting in 238 putative germination inhibitors. Secondary screens were performed on these hits to determine effects on yeast replication, cytotoxicity against mammalian fibroblasts, and direct inhibition of the NL enzyme (**Doc S1**).

To confirm that the 238 hits from the high throughput screen were bona fide inhibitors of germination, each compound was tested in the QGA. One hundred ninety-one of the 238 HTS hits showed inhibition of germination at a single, relatively high concentration (80 μM) in this assay (**Doc S2**). The majority of these confirmed germination inhibitors (121/191) showed low cytotoxicity to mammalian cells (<25% decrease in cell viability), and 167 of 191 caused at least 10% inhibition of yeast growth at 10 μM. Six of the 191 germination inhibitors were also inhibitors of the NL enzyme assay. Overall, the QGA was highly effective for validation of HTS hits and provided a high-confidence library of 191 novel germination inhibitors, resulting in the largest discovery of novel, confirmed fungal germination inhibitors in any system. Because the majority of these inhibitors exhibited low preliminary cytotoxicity against mammalian cells, the data supported the idea that germination could serve as a reservoir of fungal-specific drug targets.

Upon evaluation of the structures of the 191 confirmed inhibitors, we discovered that 76 of the compounds fell into 8 distinct groups with shared substructures, each of which contained 4 or more compounds (**Figure 2**). The identification of multiple groups of similarly structured compounds from a library of diverse small molecules is advantageous because similarly structured compounds are likely to have shared molecular targets (15,16). Compounds that had shared substructures with other inhibitors, showed potent inhibition of both spore germination and yeast growth, and exhibited low mammalian cell toxicity (**Figure 2**) were prioritized for further investigation.

**Figure 2.**
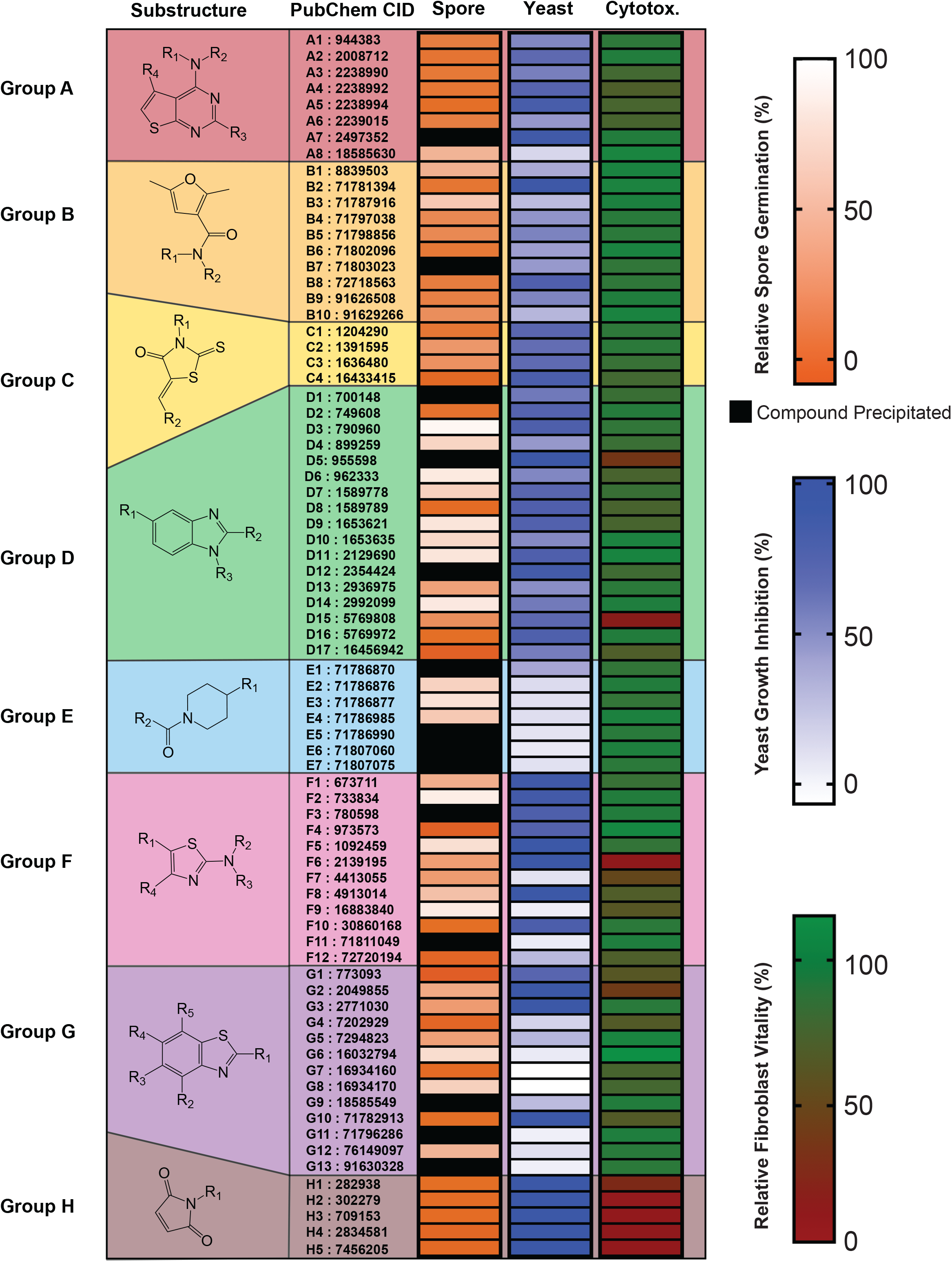
Groups (8) of compounds (76) identified and confirmed as germination inhibitors with shared substructures. Diagram of the each substructure with alphabetic assignments followed by their Pubchem CIDs for ease of identification. Heatmap representing level of relative spore germination (at 80uM), level of yeast growth inhibition (at 10uM) and level of relative fibroblast vitality (at 10uM).

#### QGA titrations of germination inhibitors identified 6 discernable phenotypes

Because our QGA data with translation inhibitors showed that inhibiting molecular targets within a specific cellular function resulted in a shared phenotype (**Figure 1**), we hypothesized that compounds with shared substructures would induce shared phenotypes. To determine potential “chemical phenotypes,” we titrated 86 confirmed germination inhibitors in the QGA. We discovered six different phenotypes, and they were distinguished from one another on the basis of differences in germination synchronicity, germination rates, and overall population behavior. These phenotypes fell into two general categories: “homogeneous germination” and “heterogeneous germination” (**Figure 3A**).

**Figure 3.**
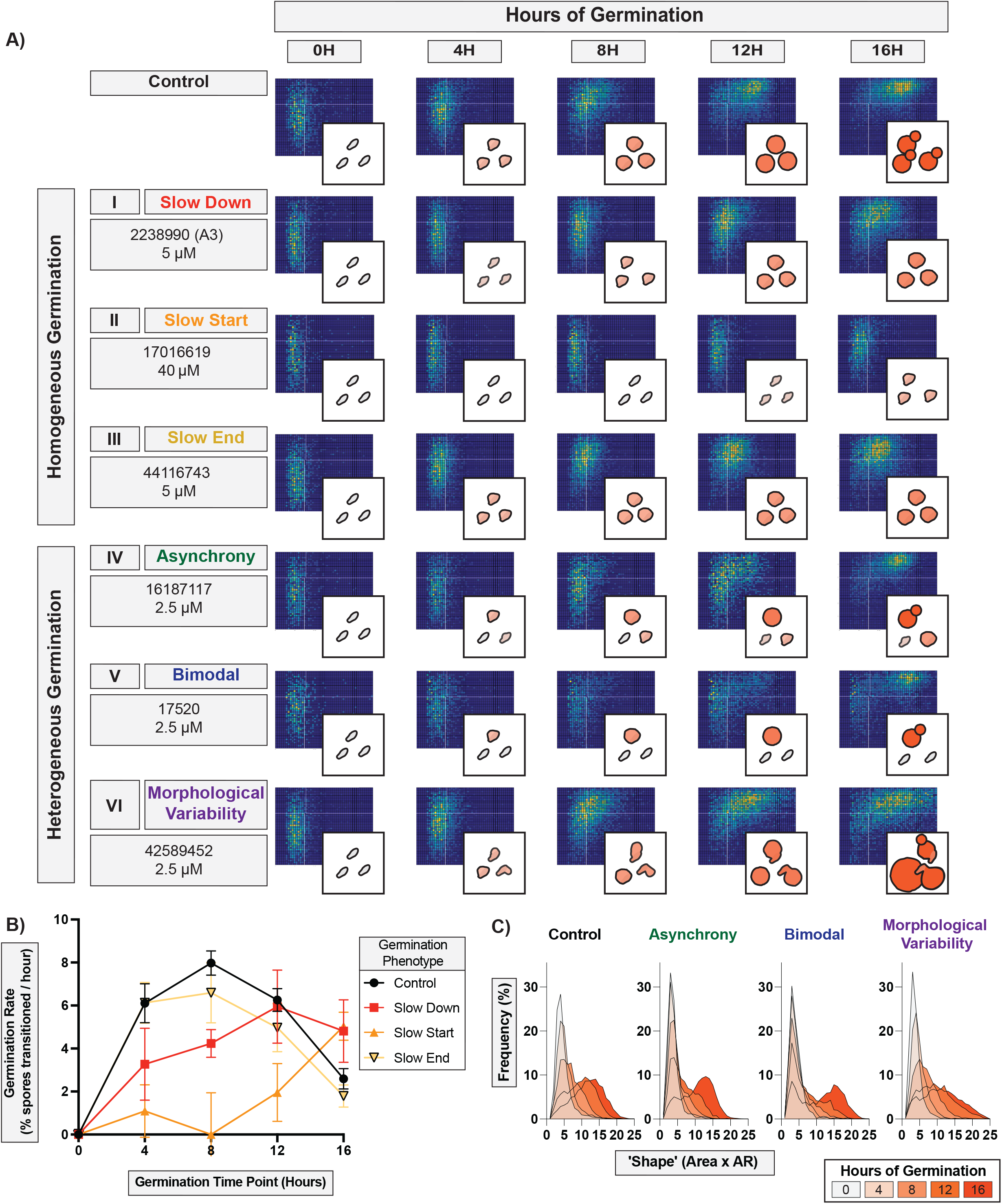
Characterization of germination inhibition phenotypes. A) Germination profiles and representative diagrams of distinct phenotypes divided into homogeneous germination (i) slow down, (ii) slow start, (iii) slow end and heterogeneous germination (IV) asynchrony, (V) bimodal, (VI) morphological variability, each plot representing ~6,000 spores B) Germination rates of spores at different time points during different homogeneous germination phenotypes (error bars represent standard deviation). C) Population morphological spread over 16 hours for heterogeneous germination phenotypes.

“Homogeneous germination” phenotypes occurred when spores germinated synchronously as a population but at a slower rate, and individual spores were affected equally throughout the population. In this category there were three phenotypes, which were distinguished by the time point at which germination was most inhibited and were quantified by changes in germination rates. The most common of these (~30% of all inhibitors tested) was the (I) Slow Down phenotype in which the population of spores germinated synchronously but at a slower rate over 16 hours of germination (**Figure 3A, rows 1-3**). The (II) Slow Start phenotype occurred with a single inhibitor at the beginning of germination. After initial inhibition (occurring between 0 and ~8 hours), the rate of germination increased rapidly as spores overcame the inhibition. The (III) Slow End phenotype occurred with a handful of inhibitors in which the population of spores initially germinated at a normal rate but exhibited a reduced rate later in germination (after ~8 hours). This inhibition was observed primarily at the point in germination (~8 hours) when spores that had germinated into small, circular cells became uniformly larger (via isotropic growth). All three of these phenotypes showed slower rates of germination, but the points of inhibition and rates of germination varied among them (**Figure 3B**).

“Heterogeneous germination” phenotypes occurred when spores no longer germinated synchronously as a population, and individual spores were affected differently. In this category, there were three phenotypes, all related to how a lack of synchrony manifested across a population. The most common of these inhibition phenotypes (~50% of inhibitors tested) was the (IV) Asynchrony phenotype, in which a population of spores lost synchronous germination, and different levels of inhibition were observed across the population, resulting in large variations in cell shapes and sizes (**Figure 3A, rows 4-6**). A (V) Bimodal phenotype occurred with several inhibitors when the population split in two groups with part of the population experiencing complete inhibition and the other germinating normally, resulting in a bimodal distribution. The (VI) Morphological Variability phenotype occurred with only two inhibitors when spores germinated into cells with a variety of sizes and shapes, resulting in a higher proportion of more elongated and/or very large cells. This led to a more variable morphology in yeast once fully germinated. These three phenotypes demonstrate that spores in a population can respond differently to germination conditions and be distinguished by population-level and individual morphological differences (**Figure 3C**). While the phenotypes observed were not necessarily a comprehensive accounting of all germination phenotypes, they provided an opportunity to further parse the 191 inhibitors into groups and evaluate structure-function relationships.

#### Similarly-structured compounds elicit the same germination phenotypes

To test the hypothesis that compounds with shared substructures would induce shared phenotypes, we evaluated structural groups A and B in more detail. If similarly-structured compounds showed similar phenotypes, it would support the idea that they share the same target. Groups A and B were chosen because they 1) contained a relatively large number of compounds with similar substructures (8 and 10, respectively), 2) displayed generally strong inhibition of germination at 80 μM, 3) were able to inhibit yeast growth to varying degrees, and 4) exhibited low cytotoxicity to mammalian cells.

Group A is composed of 8 compounds with a *Thieno[2,3-D]Pyrimidin-4-Amine* substructure. To identify the phenotypes of inhibition for group A, we carried out titrations of each at concentrations from 2.5 μM to 80 μM (**Figure 4A, B**). The majority of group A compounds (A1-A7) demonstrated clear “slow down” germination phenotypes; however, A8 showed the “asynchrony” phenotype. This phenotypic discrepancy suggests that A8 interacts differently with spores and may have a different cellular target. For this reason, A8 likely does not belong in this grouping of compounds and was considered an outlier during further characterization of group A. To determine the mammalian cytotoxicity of these inhibitors, dose response cytotoxicity assays were performed on fibroblasts with A1-A7 at concentrations from 2.5 μM to 80 μM (**Figure 4C**). These assays showed that Group A compounds exhibited relatively low cytotoxicity against mammalian cells at relevant inhibitory concentrations. Notably, potency of spore germination inhibition was not related to levels of mammalian cytotoxicity as exemplified by the strongest inhibitor in this group (A5) showing >80% germination inhibition at 5 μM and <20% cytotoxicity at concentrations as high as 80 μM. This suggests that structural moieties in this group could be altered to maximize antifungal activity while minimizing cytotoxicity. Together, these data support the hypothesis that similarly structured compounds demonstrate similar phenotypes of inhibition and that phenotypic characterization can identify outliers in structural groups. These results further support that group A compounds target the same biological process, have low cytotoxicity, and are promising candidates for antifungal development and target identification.

**Figure 4.**
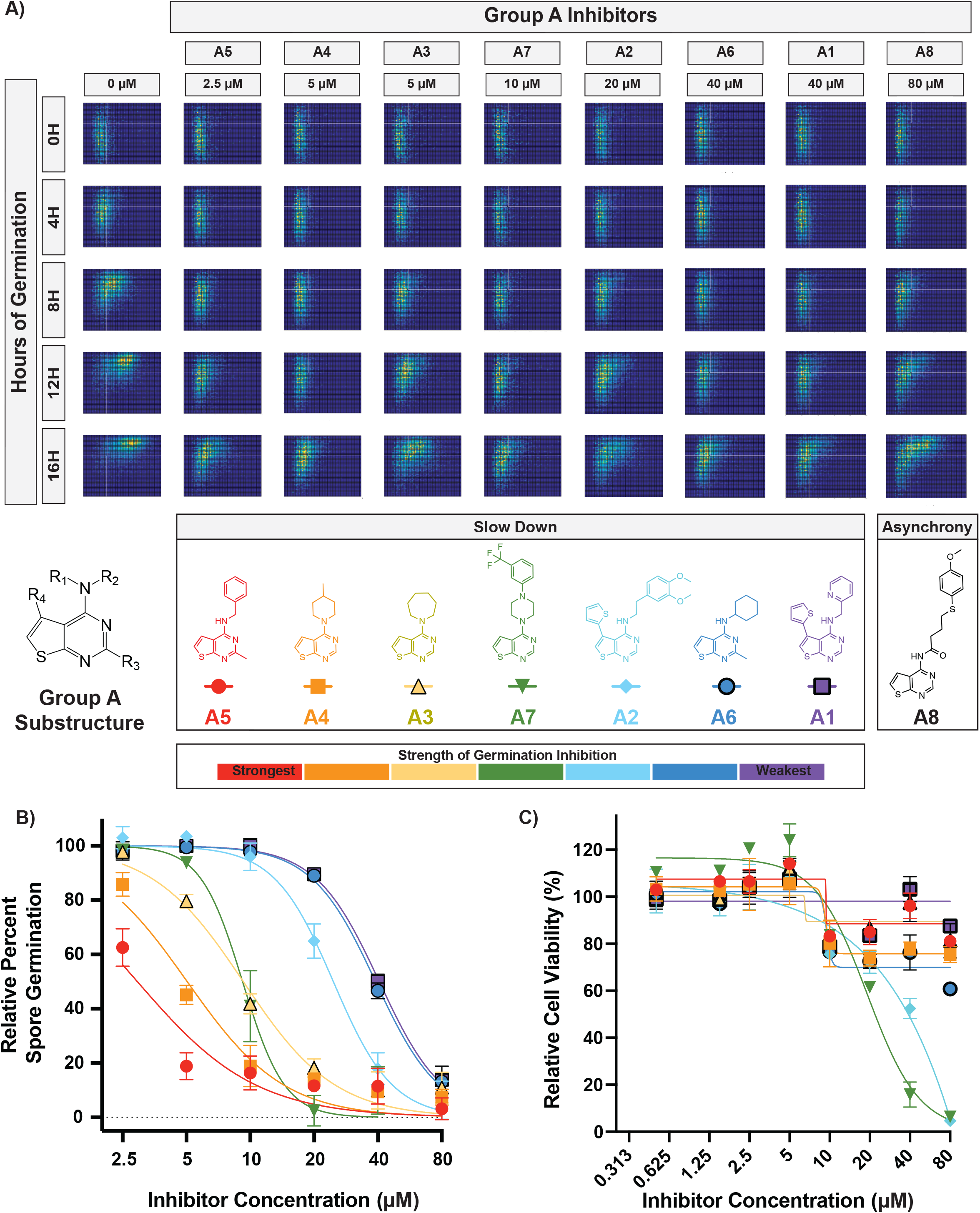
Characterization of group A compounds. Germination profiles of spores at phenotypic concentrations in the presence of group A compounds, each plot representing ~6,000 spores. B) Dose response curves of compounds A1 - A7 at concentrations from 2.5 to 80 μM (error bars represent standard deviation). If inhibitors precipitated at higher concentrations, the data point was removed. C) Cytotoxicity against mammalian fibroblasts dose response curves (error bars represents standard deviation).

Group B is composed of 10 compounds with a *2,5-dimethylfuran-3-carboxamide* substructure. To identify phenotypes of inhibition for group B, we titrated these compounds at concentrations from 2.5 μM to 80 μM (**Figure 5A, B**). While the 10 inhibitors showed varying abilities to inhibit germination, all 10 demonstrated a clear “asynchrony” germination phenotype. Cytotoxicity assays were performed at concentrations from 2.5 μM to 80 μM (**Figure 5C**) and showed that group B compounds exhibit relatively low cytotoxicity against mammalian cells at relevant inhibitory concentrations. Again, potency of spore germination inhibition was not related to levels of mammalian cytotoxicity. For example, the strongest inhibitor in this group (B2) showed >75% germination inhibition at 5 μM and ≤25% cytotoxicity at concentrations as high as 80 μM. As for Group A, these data suggest that all group B compounds target the same biological process, providing another group of promising candidates for further development.

**Figure 5.**
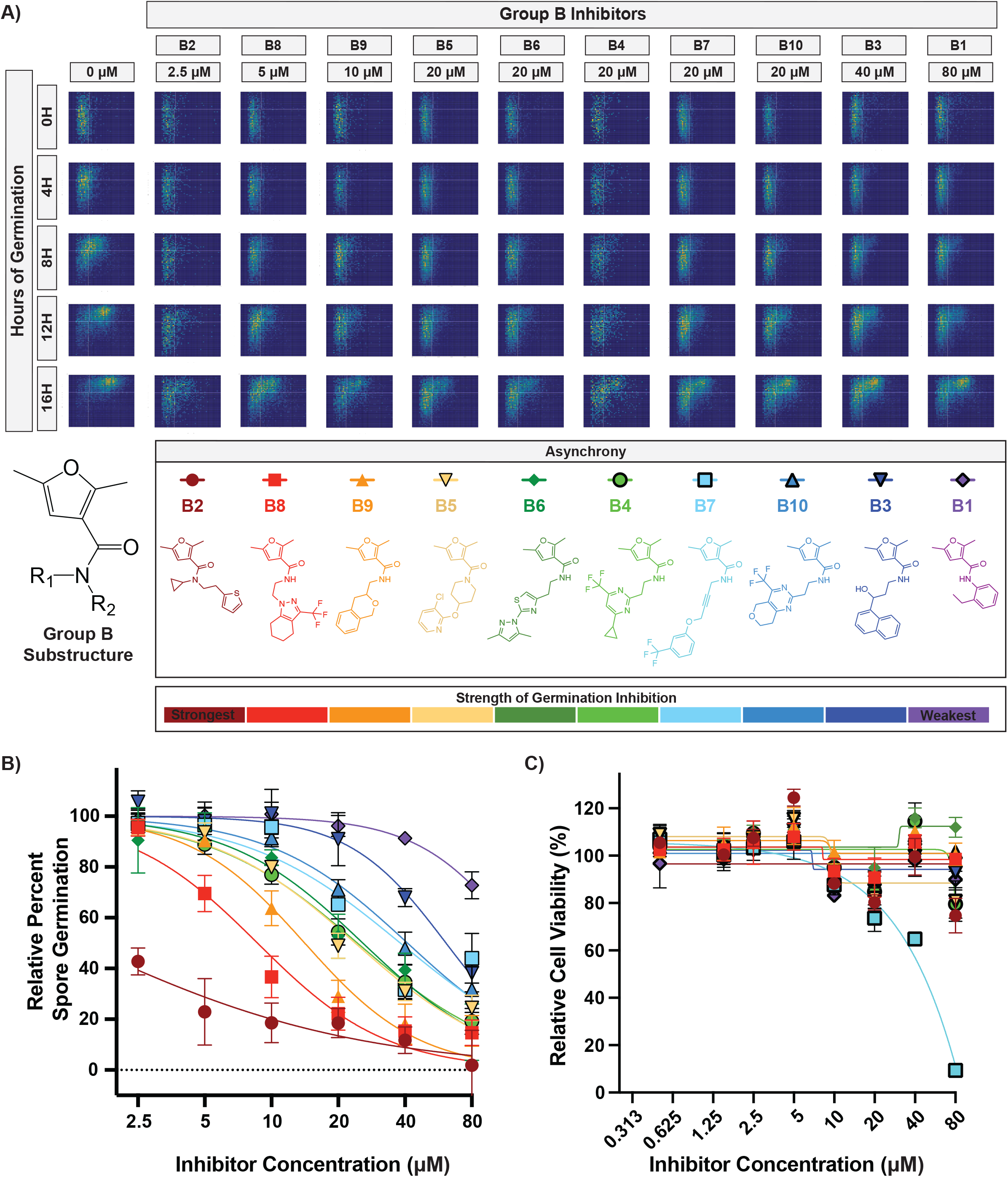
Characterization of group B compounds. A) Germination profiles of spores at phenotypic concentrations in the presence of group B compounds, each plot representing ~6,000 spores. B) Dose response curves of compounds B1 - B10 at concentrations from 2.5 to 80 μM (error bars represent standard deviation). C) Cytotoxicity against mammalian fibroblasts dose response curves (error bars represents standard deviation).

#### Complex II of the Electron Transport Chain is the likely target of Group B compounds

The relevant substructure in all the group B compounds is a *furan carboxamide*. This structure is found in known carboxamide fungicides. These fungicides have historically been used against plant pathogens and are part of the <u>S</u>uccinate <u>D</u>e<u>h</u>ydrogenase Inhibitors (SDHIs) class of fungicides, which target complex II of the electron transport chain (ETC) (17). SDHIs are a large class of fungicides that includes compounds of diverse structures that vary in their specificity for different plant fungal pathogens (17). Group B shows strong structural similarity with one SDHI in particular, furcarbanil, which has a structure nearly identical to compound B1 with only a single ethyl group differentiating the two molecules. Due to this shared similarity, we hypothesized that furcarbanil would inhibit germination at a level akin to B1 (the weakest group B compound). In fact, furcarbanil demonstrated the asynchrony phenotype at the same phenotypic concentration as B1 (80 μM), showing a nearly identical profile that was weaker than most group B compounds (**Figure 6A**).

**Figure 6.**
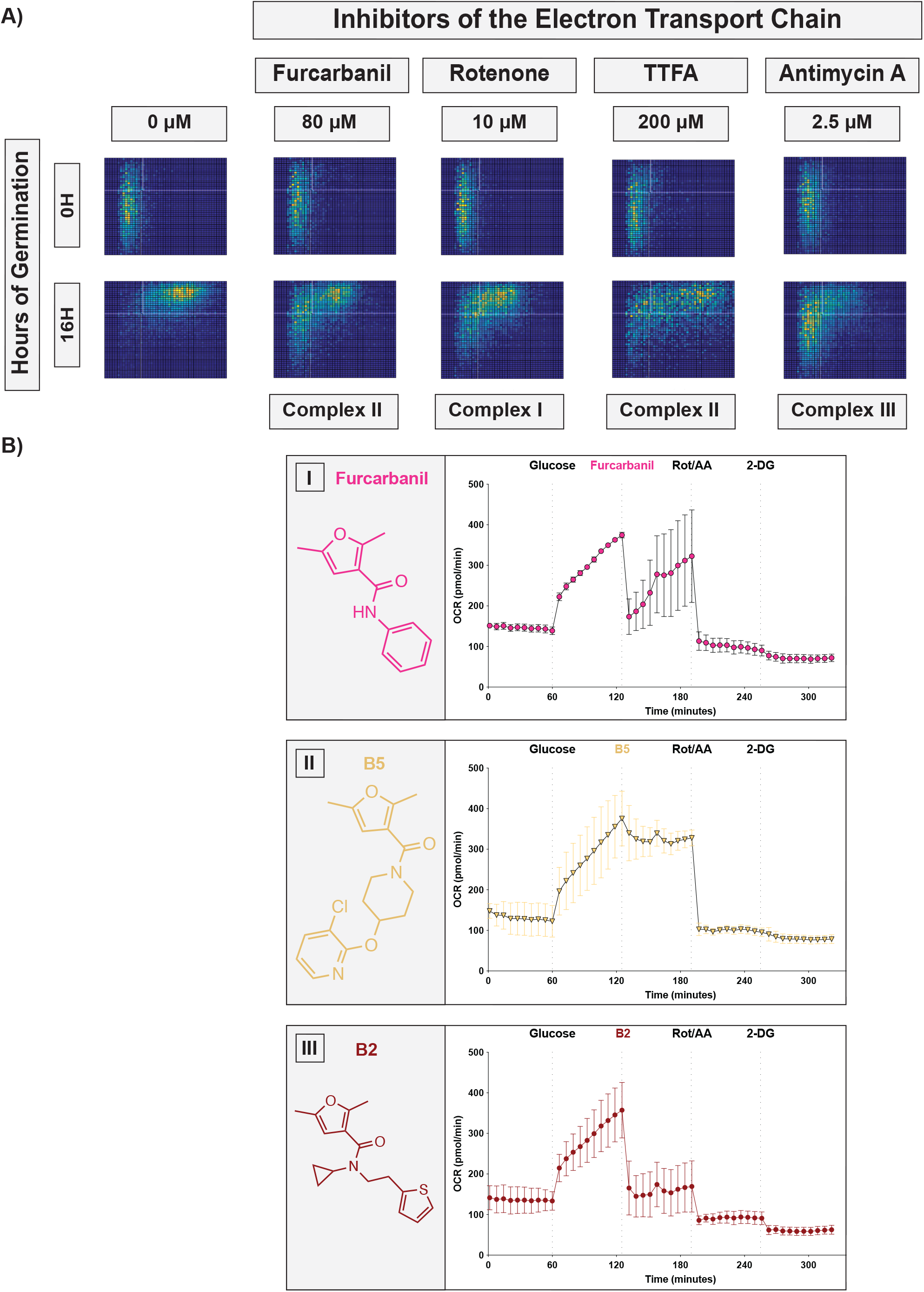
Characterization of group B compounds as electron transport chain inhibitors. A) Germination profiles of spores at phenotypic concentrations in the inhibitors Furcarbanil (80 μM), Rotenone (10 μM), TTFA (200 μM), Antimycin A (2.5 μM), each plot representing ~6,000 spores. B) Oxygen Consumption Rate (OCR) plots of JEC21 yeast with injections every 60 minutes with first Glucose (20 mM), then either (I) Furcarbanil (10 μM), (II) B5 (10 μM), or (III) B2 (10 μM), then Rotenone/AntimycinA (50 μM), and finally 2-DG (100 mM) (error bars represent standard deviation).

Given these data, we hypothesized that other inhibitors of the ETC would result in the same germination phenotype as group B compounds and furcarbanil. To test this, we determined the inhibition phenotypes of Rotenone, TTFA, and Antimycin A, which are well characterized inhibitors of Complexes I, II, and III, respectively (**Figure 6A**). Each of these mitochondrial inhibitors demonstrated the asynchrony phenotype. Given the shared structural homology of group B to furan carboxamide SDHIs and the inhibitory phenotype of furcarbanil and other ETC inhibitors, we hypothesized that group B compounds hinder germination through ETC inhibition by targeting succinate dehydrogenase (complex II) specifically. To test the possibility that group B compounds inhibit the ETC, we performed oxygen consumption experiments to monitor the effects of these inhibitors on oxygen consumption rates of *Cryptococcus* yeast (**Figure 6B**). We tested a weak (Furcarbanil), an intermediate (B5) and a strong (B1) inhibitor of germination and found that all three compounds (at 10 μM) were able to lower the oxygen consumption rate (OCR) of the cells. Furcarbanil showed initial inhibition followed by recovery, B5 showed low inhibition that was not recovered from, and B1 showed strong OCR inhibition. These results support the idea that group B compounds inhibit germination by obstructing the ETC. While it is formally possible that these inhibitors could be altering OCR through pathways other than direct ETC inhibition, all the data presented here support the hypothesis that group B compounds are novel ETC inhibitors, likely targeting complex II. These novel inhibitors exhibit strong antifungal activity and low mammalian cytotoxicity, making them prime candidates for development into novel antifungal therapeutics.

## Discussion

In this study we combined two new phenotypic assays that target fungal spore germination to identify, validate, and characterize 191 novel fungal germination inhibitors. Using QGAs, we identified 6 distinct chemical phenotypes distinguished from one another on the basis of differences in germination synchronicity, germination rates, and overall population behavior. Compounds that targeted the same cellular function or had shared substructures induced similar phenotypes. Thus, QGAs identified phenotypic outliers and facilitated target identification via comparisons between structurally similar compounds with the same phenotypes to compounds with known cellular targets. We identified a group of novel putative fungal-specific electron transport chain inhibitors that are promising candidates for antifungal development. Most importantly, this study supports the idea that the germination process holds fungal-specific pathways that could serve as targets for new antifungal drugs.

### Chemical phenotyping can help overcome the hurdles of phenotypic drug discovery

The relative merits of targeted drug discovery vs. phenotypic drug discovery have been debated across fields; however, in antifungal drug discovery, one of the key issues is the lack of known fungal specific targets. This challenge supports using phenotypic drug discovery (PDD) approaches, but PDD presents other challenges such as 1) difficulties in validation of hits, 2) an inability to establish structure-activity relationships, and 3) difficulties in target identification (18). To overcome these limitations, new methods of phenotypic characterization have been used such as molecular phenotyping in which transcriptome analysis was used as a secondary screening method (19). This approach facilitated clustering of compounds based on shared profiles and helped identify their targets. Similarly, we used new phenotypic assays to both identify and characterize compounds and overcome the hurdles of PDD.

By using the NL-based high throughput screen for inhibitors of germination (as opposed to growth), we increased the specificity of our initial screen, reducing the number of hits and increasing the likelihood that the hits would be fungal-specific. Following the initial screen with QGAs enabled identification of bona fide inhibitors of germination, eliminated false positives from the working pool of compounds, and addressed the first major challenge in PDD (validation). QGAs were also used to address the second major PDD challenge (establishing structure-function relationships) via the generation of chemical phenotypes for each compound of interest. We showed that inhibitors that target the same biological process share the same chemical phenotype. Thus, by characterizing the phenotypes of each compound in a structural group, we established structure-activity relationships and identified phenotypic outliers that could have targets that differ from the group overall. Finally, the use of chemical phenotyping also addressed the third challenge by lowering the barriers to target identification. Population dynamics of germinating spores vary in the presence of different inhibitors, stressors, nutrients and mutations, all of which can be assessed using the QGA (7,12). These provide an opportunity for comparative analyses to facilitate target identification. By combining the testing of inhibitors with alteration of nutrients, inhibition of known targets, or creation of knockout and overexpression constructs, we can mimic, intensify, alter, or eliminate a germination phenotype and thereby identify the target processes and pathways of specific inhibitors. For example, in a previous study we found that disulfiram had a slow end phenotype (7), indicating that it inhibits a target that is important for the isotropic growth phase of germination. The more we learn about the molecular programming of spore germination, the more we can gain from this type of phenotypic analysis.

### The electron transport chain as a fungal-specific target in antifungal development

The ETC has been suggested as a good target for antifungal drug development because of the role of respiration in regulating virulence traits and the existence of fungal-specific ETC elements. However, complex II/succinate dehydrogenase (SDH) has yet to be exploited (20). SDH could be a promising target, having been implicated in virulence of some human fungal pathogens. Specifically, SDH mRNA transcripts are overrepresented in *Cryptococcus* during murine pulmonary infections (21), and the SDH inhibitor Thenoyltrifluoroacetone (TTFA) has been shown to prevent hyphal formation in *Candida albicans*, a key virulence trait (22). It is unclear whether there are fungal-specific properties of SDH, but some carboxamide SDHIs have shown narrow spectrum use against basidiomycete plant pathogens (17, 23), which suggests that fungal SDH is unique. Alternatively, some SDHIs have been shown to also inhibit Complex III, implying more complex interactions (24). Nevertheless, it is promising that group B compounds in this study show high efficacy and low cytotoxicity, supporting the idea that their target(s) harbor fungal-specific features. While it is difficult to irrefutably conclude that group B inhibitors are targeting SDH, the data provided here strongly support this hypothesis. Future studies will be needed to fully characterize the mechanism of these inhibitors, to optimize them for increased antifungal potency and reduced mammalian cytotoxicity, and to test optimized inhibitors in murine models of invasive fungal infections. Overall, this group of compounds is extremely promising for further development into antifungal drugs.

### Spore germination as a target reservoir for antifungal therapeutics

Spore germination is a process that appears to be distinct from any process in humans, making it a potential reservoir for fungal-specific targets for drug development. Here, we determined that 121 of the 191 germination inhibitors we identified showed preliminary low cytotoxicity against mammalian cells, suggesting that they may be targeting fungal-specific molecules. Additionally, both group A and B compounds showed relatively low cytotoxicity at relevant inhibitory concentrations, and germination inhibition ability was not linked to cytotoxicity. This provides the opportunity to modify their structures to maximize antifungal activity while minimizing human cytotoxicity, thus increasing the difference between the effective dose and the toxic dose, leading to a higher therapeutic index. These data support the idea that targeting spore germination will result in the identification of low toxicity antifungal drug candidates.

Targeting spore germination also provides an opportunity for prevention of fungal disease, which is an area of disease management that is under-explored in the field of human fungal pathogenesis. Spores play an important role in disease progression in the majority of invasive human fungal pathogens, and spore germination is required for spores to cause disease (8, 9). Therefore, inhibiting spore germination (in addition to the subsequent vegetative replication) could provide a unique opportunity for antifungal prophylaxis in immunocompromised individuals to prevent fatal disease. The potential role of germination inhibitors in antifungal prophylaxis has been explored previously (7), and with the identification of novel inhibitors of both germination and growth, the development of preventative therapeutics can now be pursued.

## Materials and Methods

### Strains and Strain Manipulation

*Cryptococcus neoformans* serotype D (*deneoformans)* strains JEC20, JEC21, CHY3833 and CHY3836 were handled using standard techniques and media as described previously (7, 25, 26). *Cryptococcus* spores were isolated from cultures as described previously (27). Briefly, yeast of both mating types (JEC20 and JEC21 or CHY3833 and CHY 3836) were grown on yeast-peptone-dextrose (YPD) medium for 2 days at 30°C, combined at a 1:1 ratio in 1X phosphate buffered saline (PBS), and spotted onto V8 pH 7 agar plates. Plates were incubated for 5 days at 25°C, and spots were resuspended in 75% Percoll in 1X PBS and subjected to gradient centrifugation. Spores were recovered, counted using a hemocytometer, and assessed for purity by visual inspection.

### NanoLuciferase (NL) Germination Screen

All screening of the LifeChem Libraries (Life Chemicals) was carried out with the assistance of the University of Wisconsin—Madison (UW-Madison) Small Molecule Screening Facility. All NL screening was performed as described previously (7). Briefly, CHY3833 and CHY3836, reporter strains harboring a CNK01510-NL fusion construct, were used to produce spores for library screening. Screening was carried out with 1 × 10^4^ spores incubated in 384-well screening plates in 10 μL of germination medium (0.5X YPD) for 10 h at 30°C. Cells were then incubated with 10 μL of Nano-Glo luciferase assay reagent (Promega Corporation) prepared as suggested by the manufacturer at 22°C for 10 min and then read using a Perkin-Elmer Enspire plate reader at 460 nm.

### Secondary Screens

#### NL Enzyme Test

CHY3833 was grown overnight in liquid YPD, washed 3 times in 1X PBS and resuspended to an OD_600_=1.00. Cells (100 μL) were added to 384-well plate wells containing 10 μM of each compound and were incubated with 10 μL of Nano-Glo luciferase assay reagent (Promega Corporation) at 22°C for 10 min and then read using a Perkin-Elmer Enspire plate reader at 460 nm. Compounds that caused a >50% decrease in luciferase signal were determined to be NanoLuciferase enzyme assay inhibitors.

#### Yeast Replication

CHY3833 and CHY3836 were each grown in YPD liquid overnight at 30°C to saturation and then resuspended in 0.5X YPD at an OD_600_ of 0.005. Strains aliquoted into 384 well plates with inhibitors and grown for 12 hours at 30°C at 3000 RPM before OD_600_ readings were taken. Compounds that caused a >10% decrease in growth were considered yeast growth inhibitors.

#### Fibroblast cytotoxicity

1 × 10^3^ normal human dermal fibroblasts (NHDF) cells per well were plated in a 384 well plate in cell culture medium. The cells were incubated with each compound of interest at 10 μM concentration for 72 hours at 37°C + 5% CO_2_. Following treatment, CellTiter-GLO (Promega) reagent was used to assay ATP dependent luminescence and thus provide a measure of cell viability. Compounds that resulted in <75% cell viability were considered low toxicity.

### Quantitative Germination Assay

Germination assays were modified from Barkal et al. 2016 to introduce automation, increase throughput, and refine assay consistency (12). Briefly, 384 well plates (Thermo Scientific: 142762) were loaded with 10^5^ spores per well, and at 0 h, synthetic medium + 2% dextrose (SD) medium containing the compounds of interest was added to the sample (final volume of 40 μL). All compounds were tested initially at 80 μM; however, concentrations were changed on a case-by-case basis for subsequent experiments. All assays and controls were performed with a final concentration 0.8% DMSO (the compound solvent) unless specified otherwise. Spores were germinated at 30°C in a humidified chamber, and the same ~5 × 10^3^ cells were monitored every 2 h for 16 h. Imaging was performed on a Ti2 Nikon microscope, and each condition was visualized in a minimum of two individual wells with three fields of view acquired from each well. All images were analyzed as described previously based on cell shape and size using ImageJ. The population ratios of spores, intermediates, and yeast were determined. Error bars in plots are based on the variation among all fields of view acquired. Level of germination was determined by quantifying the decrease in the proportion of spores in a population, and rates were quantified by determining the change in this proportion over time.

#### Translation inhibitors and concentrations

Cycloheximide (0.078 μM – 80μM) (Dot Scientific, Inc: DSC81040-1), G418 (1.6mM) (Fisher Scientific: AAJ6267106), and Puromycin (20 mM) (Dot Scientific, Inc: DSP33020-0.025) were tested with no DMSO (all were water-soluble), whereas Anisomycin (62.5 μM) (Sigma-Aldrich: A9789-5MG) was tested in 2.5% DMSO.

#### Mitochondrial inhibitors and concentrations

Furcarbanil (80 μM) (Sigma-Aldrich: T313122), Rotenone (10 μM) (Fisher Scientific: 501687383) and Antimycin A (2.5 μM) (Santa Cruz Biotechnology, Inc: sc-202467A) were all tested in 0.8% DMSO, and TTFA (200 μM) (Sigma-Aldrich: 88300-5G) was tested in 3% DMSO.

### Oxygen Consumption Rate (OCR) Experiments

Oxygen consumption experiments were performed on a Seahorse Biosciences XFe96 Extracellular Flux Analyzer, and the assays were modified from Lev et al 2020 (28). Briefly, cartridges were hydrated overnight in Agilent Seahorse XF calibrant. JEC21 yeast were grown overnight in YPD, washed with ddH_2_0 and resuspended to OD_600_=0.8 in pH 7.4 (Agilent: 103575-100). Cells (180 μL) were loaded into each well with the exception of blank wells, which were filled with XF DMEM Medium. Each injection solution was 10X the final volume in XF DMEM Medium. The assay was carried out at 30°C with injections occurring every 60 minutes. OCR was read every 6 minutes during each hour interval with 3 minutes of mixing and 3 minutes of measuring. The following final concentrations were achieved after each injection: a) 20 mM dextrose b) 10 μM of chosen inhibitor (furcarbanil, B2, or B5) and 1%DMSO c) 50 μM Rotenone, 50 μM Antimycin A and 1% DMSO and d) 100 mM 2-DG (Sigma-Aldrich: D8375-1G). Each condition was tested in triplicate.

## Acknowledgments

We thank the staff at the Small Molecule Screening Facility at the University of Wisconsin Cancer Carbone Center for their assistance and expertise with support from National Institutes of Health (NIH) grant P30 CA014520. We specifically thank Gene Aniev, Spencer Ericksen, Song Guo, and Scott Wildman for their assistance and input. We also thank Hunter Gage, Megan McKeon, Anna Frerichs and Eddie Dominguez lab for their comments on the manuscript. Finally, we thank James (Muse) Davis for his microscopy brilliance in helping to automate our germination assay.

This study was supported by an Individual Biomedical Research Award from The Hartwell Foundation to C.M.H., an NIH grant R01 AI137409 to C.M.H., and an HHMI Gilliam Fellowship to S.C.O.

---

**Figure S1.**
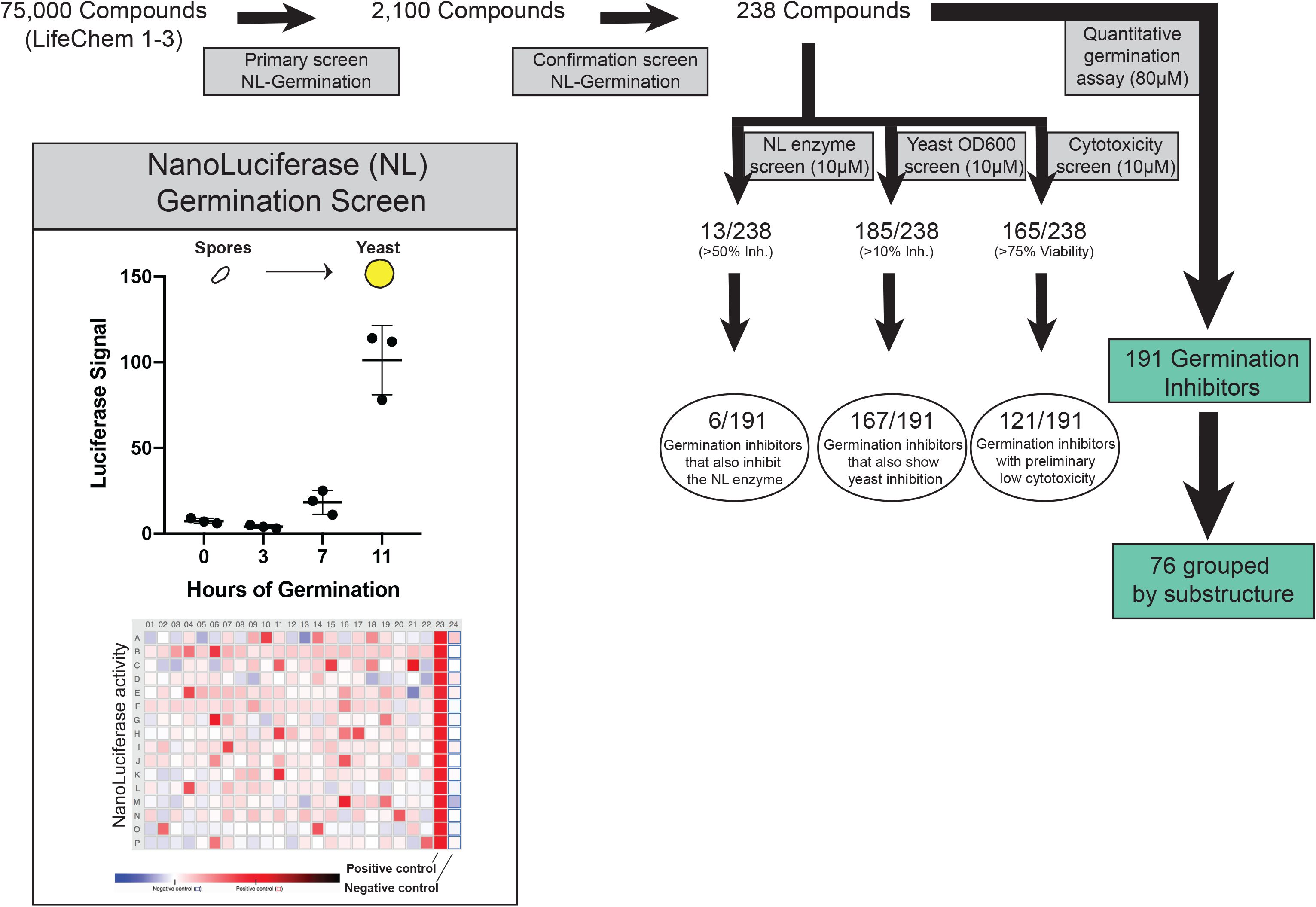
Flow chart of screening process. Box (top): spore containing NanoLuciferase (NL) construct demonstrate an increase in NL signal over the course of germination, x-axis is hours of germination and y-axis is Luciferase signal (error bars represent standard deviation among replicates). Box (bottom): representation of a 384 well screening plate. The positive control for inhibition (column 23) is 3.55 μM cycloheximide and the negative control (column 24) is 0.1% DMSO.

